# Recapitulating evolutionary divergence in a single *cis*-regulatory element is sufficient to cause expression changes of the lens gene *Tdrd7*

**DOI:** 10.1101/2020.03.22.002535

**Authors:** Juliana G. Roscito, Kaushikaram Subramanian, Ronald Naumann, Mihail Sarov, Anna Shevchenko, Aliona Bogdanova, Thomas Kurth, Leo Foerster, Moritz Kreysing, Michael Hiller

## Abstract

Mutations in *cis*-regulatory elements play important roles for phenotypic changes during evolution. Eye degeneration in the blind mole rat (BMR) and other subterranean mammals is significantly associated with widespread divergence of eye regulatory elements, but the effect of these regulatory mutations on eye development and function has not been explored. Here, we investigate the effect of mutations observed in the BMR sequence of a conserved non-coding element upstream of *Tdrd7*, a pleiotropic gene required for lens development and spermatogenesis. We first show that this conserved element is a transcriptional repressor in lens cells and that the BMR sequence partially lost repressor activity. Next, we recapitulated the evolutionary changes by precisely replacing the endogenous regulatory element in a mouse line by the orthologous BMR sequence with CRISPR-Cas9. Strikingly, this repressor element has a large effect, causing a more than two-fold up-regulation of *Tdrd7* in developing lens. Interestingly, the increased mRNA level does not result in a corresponding increase in TDRD7 protein nor an obvious lens phenotype, likely explained by buffering at the posttranscriptional level. Our results are consistent with eye degeneration in subterranean mammals having a polygenic basis where many small-effect mutations in different eye-regulatory elements collectively contribute to phenotypic differences.

## INTRODUCTION

Colonization of new habitats, such as the underground environment, is often linked to morphological, physiological and behavioral adaptations that confer advantages in the new habitat. Among mammals, the naked mole rat, blind mole rat, star-nosed mole, and cape golden mole comprise four distinct lineages that have independently evolved adaptations to live in such constantly dark (or poorly illuminated) environments. One of the most striking adaptations of these subterranean mammals is the reduction or loss of the visual system, evident by the presence of a degenerated lens and retina, as well as a reduction of the visual-processing area of the brain (1–8).

Genetically, the degeneration of the visual system is related to mutations in eye-related genes. Previous studies that investigated the genomes of subterranean mammals showed that many genes involved in eye development and function are diverged or lost in these species (9–14). Naturally-occurring or laboratory-induced gene-inactivating mutations in some of these genes in humans or mice cause malformation of eye structures and impaired vision (15–19), suggesting that the gene losses observed in subterranean mammals contributed to the evolution of degenerated eyes.

In addition to the widespread loss of genes, recent studies discovered divergence in many eye regulatory elements (20–23). In particular, genome-wide analysis of conserved non-coding elements (CNEs) revealed that CNEs with preferential sequence divergence in subterranean mammals are significantly associated with eye-related genes and significantly overlap regulatory elements that are active in developing and adult eyes of mice or humans (21). Sequence divergence in these CNEs also resulted in a large-scale divergence of binding sites of eye-related transcription factors (21–23). Together, this suggests that widespread divergence in both genes and in *cis*-regulatory elements may have contributed to the degeneration of eye structures in subterranean mammals, leading to a greatly impaired visual system.

In protein-coding genes, premature stop codon, frameshift or other reading frame-inactivating mutations can be considered as equivalent if they result in a non-functional protein. This makes it possible to use phenotypes observed in mouse knockout lines or human individuals with inactivated genes to infer the effect of evolutionary gene losses, even though the identity of the underlying inactivating mutations is different. In contrast to genes, where inactivating mutations are often predictable, the effects of mutations in *cis*-regulatory elements is less clear and mostly unexplored. In particular, it is hard to predict how, and to what extent, mutations in regulatory elements affect its activity and how such activity changes in turn affect the expression of its target gene. Consequently, to understand the effect of regulatory mutations *in vivo*, it is necessary to precisely recapitulate the naturally-occurring mutations on a model organism and quantify the consequences at the molecular and morphological level. Considering the importance of *cis*-regulatory changes for morphological evolution (24,25), characterizing the effect of naturally-occurring mutations in regulatory elements *in vivo* is crucial to understand the evolution of phenotypic traits.

To explore the effect of regulatory divergence on eye degeneration in subterranean mammals, we focused on a lens regulatory element near *Tdrd7* (tudor domain-containing 7), a gene essential for lens development and spermatogenesis. We used CRISPR-Cas9 to precisely replace the mouse sequence of this regulatory element by the blind mole rat sequence and characterized the molecular and morphological effects of the mutations observed in the blind mole rat on lens development and function.

## RESULTS

### Divergence in a lens regulatory element near Tdrd7

In previous studies, we used two comparative genomics approaches to identify and associate sequence and transcription factor (TF) binding site divergence in CNEs to the vision-impaired phenotype of the subterranean blind mole rat, naked mole rat, cape golden mole, and star-nosed mole (21,22). Both screens detected significantly higher sequence and binding site divergence in subterranean mammals in a locus that comprises two CNEs located 97 bp from each other (Figures 1A,B). These CNEs are part of a larger 339 bp region that evolves under constraint according to GERP++ (26). Higher sequence divergence is evident in the star-nosed mole, cape golden mole and, most prominently, in the blind mole rat, which exhibits a 197 bp deletion (Figures 1A,B).

**Figure 1.**
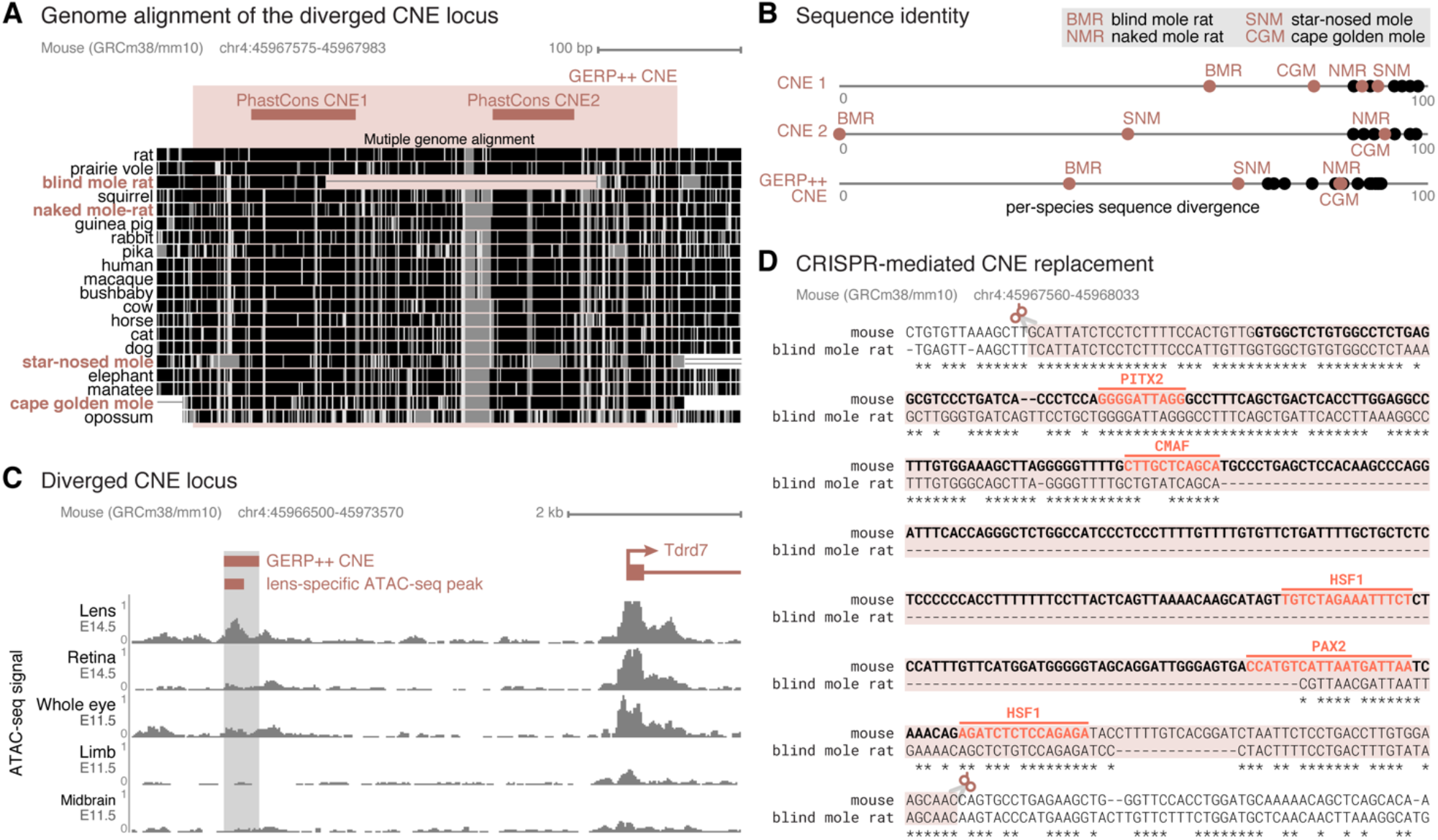
Divergence in a lens regulatory element near *Tdrd7* in subterranean mammals. (A) Multiple genome alignment, showing the two diverged CNEs detected with PhastCons and the larger CNE detected with GERP++ that encompasses both PhastCons CNEs. In the multiple genome alignment, aligning sequence is visualized by black and grey boxes. The darker the color of the box, the higher is the sequence similarity in the alignment. The subterranean vision-impaired species are highlighted in red. (B) Sequence divergence plot of the conserved elements shown in (A). The x-axis shows the percent identity between the CNE sequence of a species and the reconstructed placental mammal ancestor. Subterranean mammals are represented by red dots, all other mammals by black dots. The star-nosed mole, cape golden mole and, most prominently, the blind mole rat exhibit more diverged CNE(s) compared to other mammals. (C) Overview of the larger genomic context, showing the GERP++ CNE, the *Tdrd7* gene located ~4.5Kb downstream and ATAC-seq signal tracks of different developing mouse tissues. The CNE overlaps an ATAC-seq peak observed in lens of E14.5 mice. (D) Alignment between mouse and blind mole rat sequence encompassing the GERP++ CNE. The mouse CNE sequence is in bold. The sequence highlighted by the red box corresponds to the genomic mouse sequence that was replaced by the orthologous blind mole rat sequence with CRISPR-Cas9. Predicted transcription factor binding motifs are highlighted in the mouse sequence.

In mouse, this larger CNE overlaps many epigenetic marks derived from embryonic and adult mouse eye tissues (Figure 1C). In particular, we observe ChIP-seq peaks for eye transcription factors such as OTX2, CRX and NRL (27–29). In addition, the mouse sequence has predicted binding sites for TFs relevant for eye development, such as PITX2, CMAF, HSF1 and PAX2, three of those sites overlapping the large blind mole rat deletion (Figure 1D). Furthermore, this region overlaps ATAC-seq and DNAseI hypersensitivity peaks in adult eye tissues (30–32) as well as a prominent lens-specific ATAC-seq peak in developing mouse lens (21) (Figure 1C). Together, this indicates that the CNE may function as a regulatory element in lens tissue.

This CNE is located ~4.5 kb upstream of the conserved transcription start site of *Tdrd7* (Figure 1C), a gene that is highly expressed in lens and testis (33–35). The encoded TDRD7 protein is a component of cytoplasmic ribonucleoprotein granules that are involved in the posttranscriptional control of genes critical for lens development and spermatogenesis (33,35). Mouse studies showed that knock-down or loss-of-function mutations in *Tdrd7* lead to lens malformation, cataracts and glaucoma (33). In addition, male *Tdrd7* knockout mice are sterile due to an arrest in spermatogenesis (35). Human individuals carrying *TDRD7* loss-of-function mutations also exhibit cataracts and azoospermia (33,36). Thus, *Tdrd7* is essential for normal lens development and function and spermatogenesis. Importantly, in contrast to divergence of the CNE sequence, *Tdrd7* is intact in all subterranean mammals, including the blind mole rat, likely because of its pleiotropic role in lens development and fertility.

The close proximity to a gene required for normal lens development, and the overlap with several epigenetic marks identified in different eye tissues suggest that the genomic locus containing the diverged CNE may function as a regulatory element controlling *Tdrd7* expression. Our observation that this CNE is highly diverged in a subterranean mammal with degenerated lenses makes this CNE a promising candidate to test whether the naturally-occurring mutations have an effect on *Tdrd7* expression and normal development or function of the lens.

### The *Tdrd7* CNE acts as a transcriptional repressor in mouse lens cells

We first tested the mouse and blind mole rat (BMR) sequences for regulatory activity, using an *in vitro* dual luciferase reporter gene assay in a lens cell line. We cloned both sequences in the same genomic orientation with respect to the *Tdrd7* promoter into a firefly luciferase vector, co-transfected this vector with a control renilla luciferase vector into lens cells, and measured firefly luciferase expression driven by the mouse and BMR sequences. This experiment revealed that the mouse sequence is indeed a regulatory element (Figure 2). Interestingly, the mouse sequence acts as a repressor element, as firefly luciferase expression was significantly below the expression obtained with a control vector that does not contain any sequence insert upstream of the promoter. The expression driven by the BMR sequence was also lower than the control vector expression, but was significantly higher than the mouse sequence. This observation suggests that sequence changes, including the larger deletion observed in the BMR sequence, result in a partial release of repressor activity.

**Figure 2.**
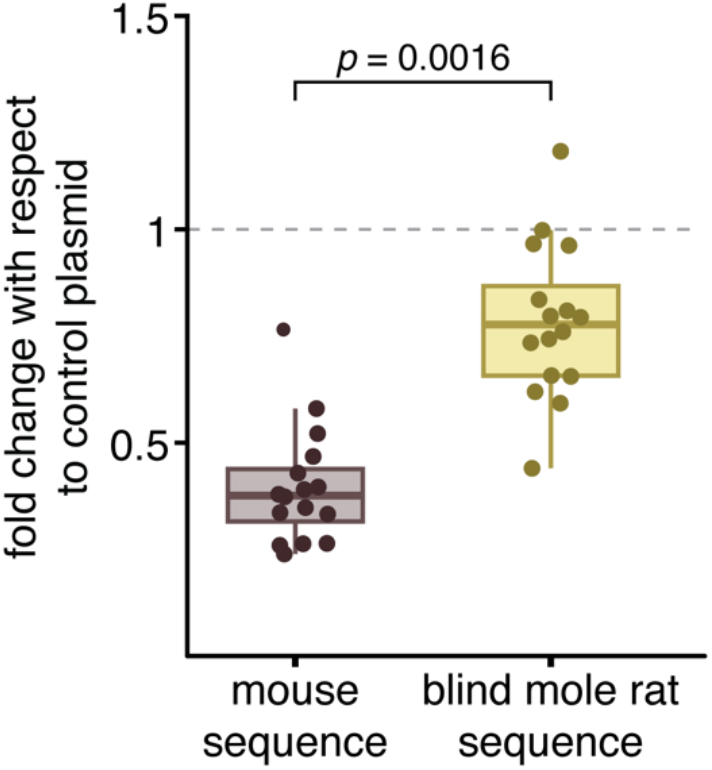
Partial loss of repressor activity of the blind mole rat CNE sequence. *In vitro* dual luciferase assay shows that the blind mole rat sequence (yellow) of the *Tdrd7* CNE exhibits a significantly weaker repressor activity than the mouse sequence (purple). Expression values correspond to the fold change with respect to the control plasmid that lacks any sequence insert upstream of the promoter. The *p*-value was computed with a two-sided Wilcoxon rank sum test.

### Recapitulating evolutionary changes in the lens regulatory element in mouse

The observed partial loss of repressor activity of the BMR sequence predicts that replacing the mouse genomic sequence of this regulatory element by the BMR sequence may result in an up-regulation of *Tdrd7*. To test this *in vivo*, we created a transgenic mouse line (referred hereafter as BMR^*repr*^ mice) in which we recapitulated the naturally-occurring BMR mutations in this repressor element. To this end, we used CRISPR-Cas9 to precisely replace the 410 bp endogenous mouse sequence encompassing the lens regulatory element with the orthologous BMR sequence (Figure 1D). We used C57BL/6J mice because the N strain already carries a mutation in the photoreceptor *Crb1* gene, resulting in retinal degeneration (37). BMR^*repr*^ mice were born at normal Mendelian ratios, were viable and fertile and showed no obvious phenotype. In the following, we used the BMR^*repr*^ mouse line to quantify the molecular and morphological effects of evolutionary divergence in this lens regulatory element.

### *Tdrd7* is up-regulated in the developing lens of BMR^*repr*^ mice

To validate the partial loss of lens repressor activity of the BMR sequence *in vivo* and to test whether the observed repressor activity difference affects *Tdrd7* expression in the lens, we used RT-qPCR to compare the expression of *Tdrd7* in wild-type and homozygous BMR^*repr*^ mice at three different time points, embryonic day 14.5 (E14.5), postnatal day 1 (P1) and three months (adult). Since *Tdrd7* has an additional role in spermatogenesis, we also quantified its expression in the testis and ovaries at the same time points.

We found that *Tdrd7* is consistently and significantly up-regulated in the lens of E14.5 and P1 BMR^*repr*^ mice in comparison to wild-type mice (Figure 3A). No differences in *Tdrd7* expression were observed in adult lens. We also observed a small but marginally-significant down-regulation of *Tdrd7* in adult but not developing testes, and a marginally-significant up-regulation in developing ovaries (Figure 3B). Despite *Tdrd7* expression changes in reproductive organs, increased *Tdrd7* expression is consistently observed in developing lenses of male and female mice, and the magnitude of *Tdrd7* up-regulation is substantially higher in the lens compared to the magnitude of *Tdrd7* expression differences in testis and ovary (Figures 3A, B). Furthermore, the absolute expression level in ovaries of wild-type and BMR^*repr*^ mice is substantially lower than in the lens (Figures 3C,D). Together with the ATAC-seq data and luciferase reporter assays, this suggests that the major function of the regulatory element is to suppress *Tdrd7* expression in the developing lens, which reveals an unknown aspect of *Tdrd7* regulation.

**Figure 3.**
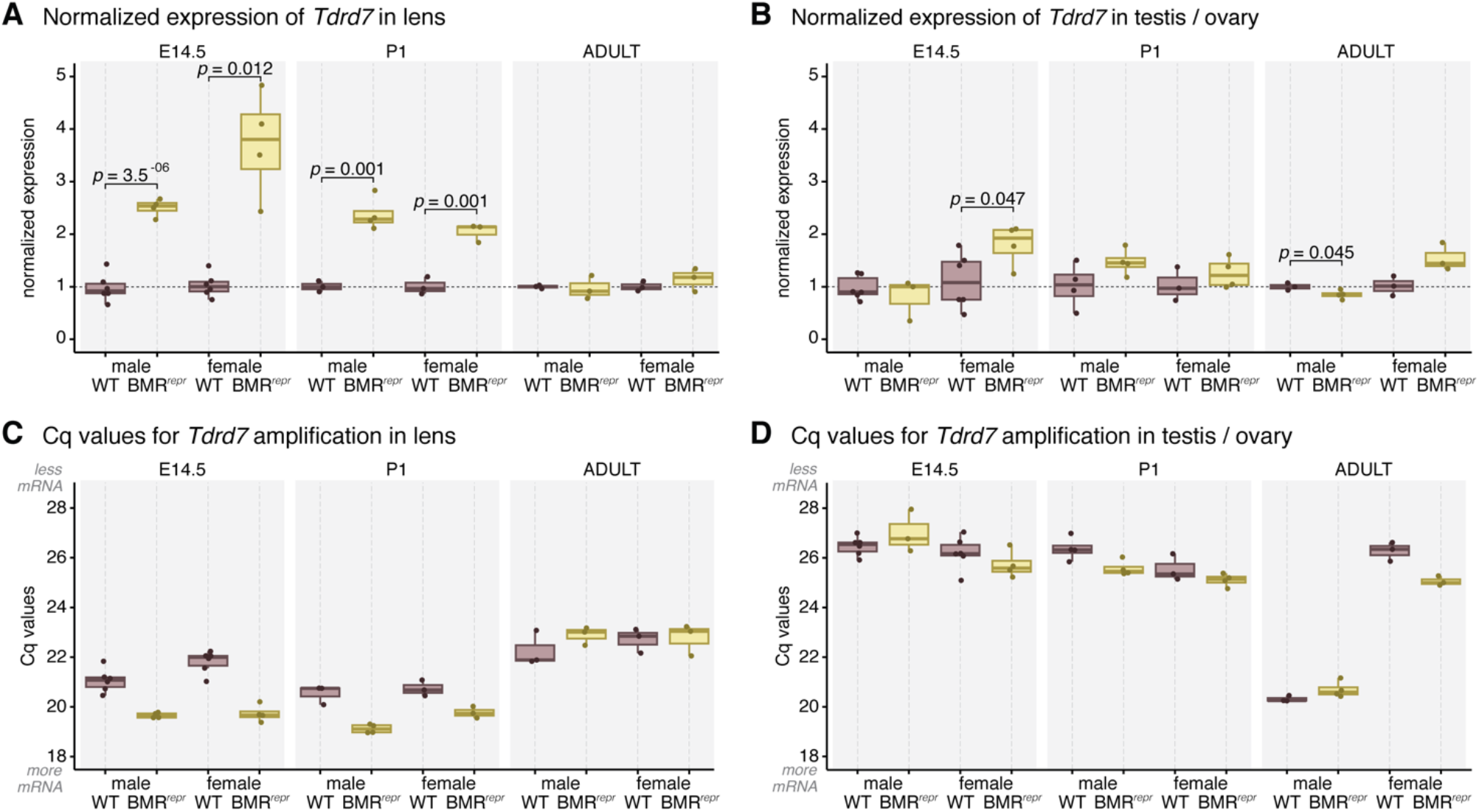
*Tdrd7* is up-regulated in developing lens of BMR^*repr*^ mice. (A, B) Comparison of *Tdrd7* expression in developing and adult lens (A) and reproductive organs (B) of wild-type (purple) and BMR^*repr*^ mice (yellow) with RT-qPCR. The strongest difference is a more than two-fold up-regulation of *Tdrd7* in developing lens of BMR^*repr*^ mice. Expression values were normalized by the housekeeping gene *Rpl13a* and by the levels of wild-type mice. Each data point is a biological replicate. *P*-values were computed with a two-sided t-test. (C, D) Comparison of Cq values indicate that *Tdrd7* mRNA is more abundant in the lens and adult testis than in the developing testis or ovary.

### BMR^*repr*^ mice have no obvious lens phenotype

Knock-down and loss-of-function mutations in *Tdrd7* have been shown to impair normal lens morphology and function (33,35), but the morphological effects of an up-regulation of *Tdrd7* have not yet been investigated. Therefore, we analyzed whether the observed increase in *Tdrd7* expression in the developing lens of homozygous BMR^*repr*^ mice affects lens function and morphology. We first investigated whether lenses of adult BMR^*repr*^ mice exhibit signs of cataracts, a clouding of parts of the lens caused by aggregation of lens proteins (38). To this end, we carefully dissected lenses of wild-type and BMR^*repr*^ mice, imaged them under darkfield illumination and quantified opaque cataract aggregates in the core of the lens, which is the only place where we observed such aggregates (Figure 4A). As shown in Figure 4B, we found that BMR^*repr*^ mice have a slightly higher number of cataract aggregates compared to wild-type mice (average 6.06 vs. 4.47; Wilcoxon two-sided rank test *p*-value of 0.18). The size of cataract aggregates is slightly lower in BMR^*repr*^ compared to wild-type mice (average 14.2 μm vs. 15.5 μm; Wilcoxon two-sided rank test *p*-value of 0.33). To test whether the subtle difference in cataract aggregate number is augmented in older BMR^*repr*^ mice, we analyzed individuals older than 1.5 years, but found no increase (Figure 4B).

**Figure 4.**
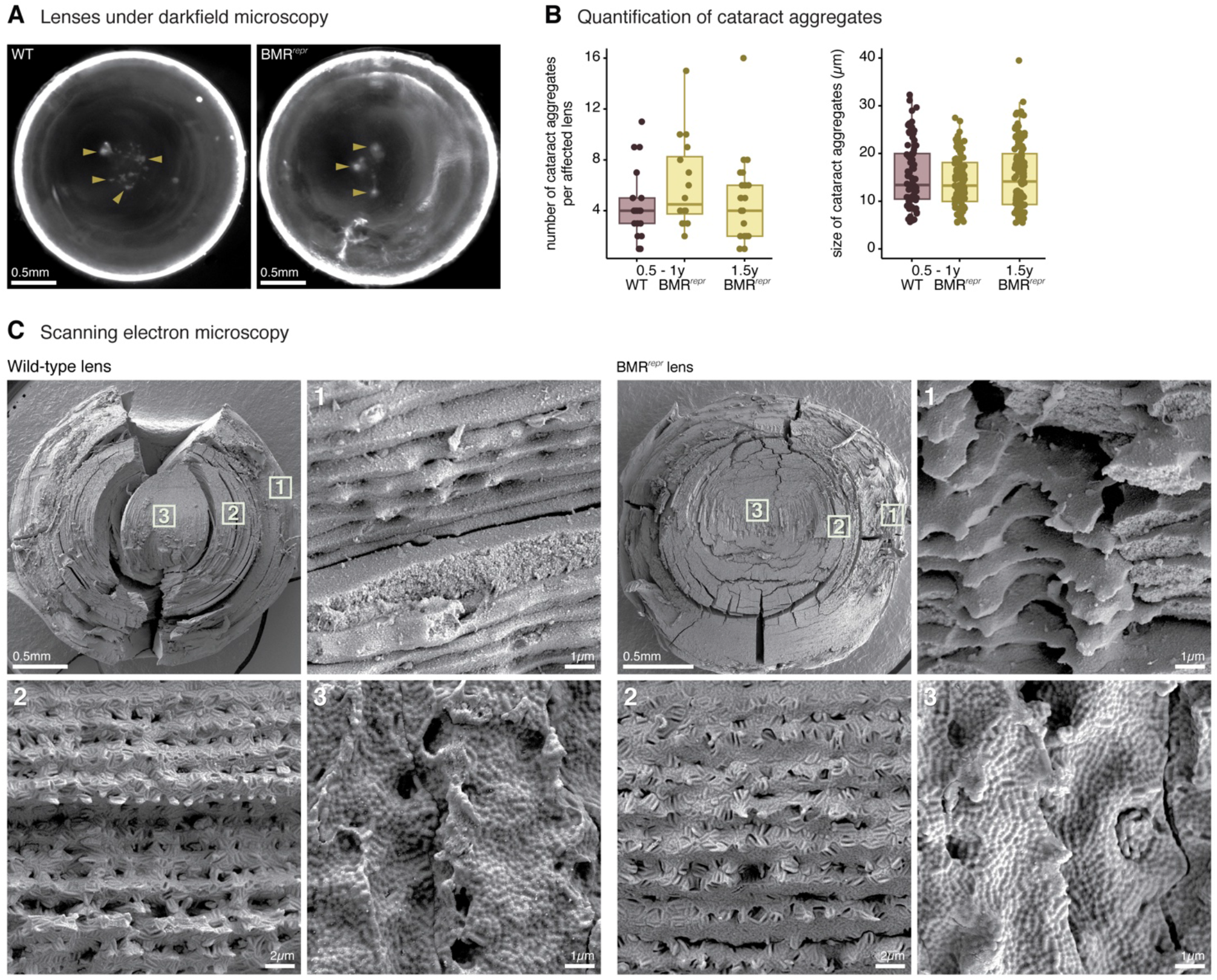
BMR^*repr*^ mice do not show an obvious lens phenotype. (A) Lenses of wild-type and BMR^*repr*^ mice viewed under darkfield illumination exemplify cataract aggregates (yellow arrowheads) in the center. (B) Quantification of the number and size of cataract aggregates in age-matched (0.5 to 1 year old) wild-type (purple) and BMR^*repr*^ (yellow) mice shows that BMR^*repr*^ mice have a slightly higher number of similarly-sized cataract aggregates (differences are not statistically significant). We additionally analyzed 24 lenses of 1.5-year old BMR^*repr*^ mice and these showed no increase in number or size of cataract aggregates compared to younger BMR^*repr*^ mice. (C) Comparison of lens fibers in 2 year old wild-type and BMR^*repr*^ mice with scanning electron microscopy shows no obvious morphological differences. Different areas of the lens, highlighted by boxes with numbers 1-3 at the top left most image, are represented in closer detail in the remaining images.

Next, we investigated whether lens fibers of BMR^*repr*^ mice exhibit structural changes compared to those of wild-type mice. The lens is composed of multiple concentric layers of tightly-connected fibers. The fiber membranes have numerous and complex interdigitations and cell-cell junctions, resulting in a rigidly structured epithelium that is important for lens function by minimizing the spacing between fibers and, thus, minimizing light scattering (38). We used scanning electron microscopy to image the morphology of the lens fibers of adult mice and compared the outer, median, and inner layers of the lenses. As shown in Figure 4C, there is no obvious difference in shape or fiber arrangement between wild-type and BMR^*repr*^ mice.

Together, these analyses show that the substantial increase in *Tdrd7* expression in the developing lens of BMR^*repr*^ does not result in a clearly-detectable lens phenotype.

### Up-regulation of *Tdrd7* does not result in an increased TDRD7 protein content

Given the lack of an obvious lens-related phenotype, we investigated whether the significantly increased *Tdrd7* mRNA levels in the lens of BMR^*repr*^ mice lead to an increase in TDRD7 protein levels. To this end, we used mass spectrometry to precisely quantify the absolute amount of TDRD7 in the lens of wild-type and homozygous BMR^*repr*^ mice at E14.5 and P1, the two time points where *Tdrd7* mRNA expression is significantly increased. Surprisingly, we observed a similar amount of TDRD7 protein in the lens of wild-type and BMR^*repr*^ mice (Figure 5). This indicates that differences in *Tdrd7* expression are effectively buffered at the protein level by a currently unknown mechanism. The absence of significantly different protein levels provides an explanation of why the embryonic mis-regulation of *Tdrd7* in the lens of BMR^*repr*^ mice does not result in a detectable lens phenotype.

**Figure 5.**
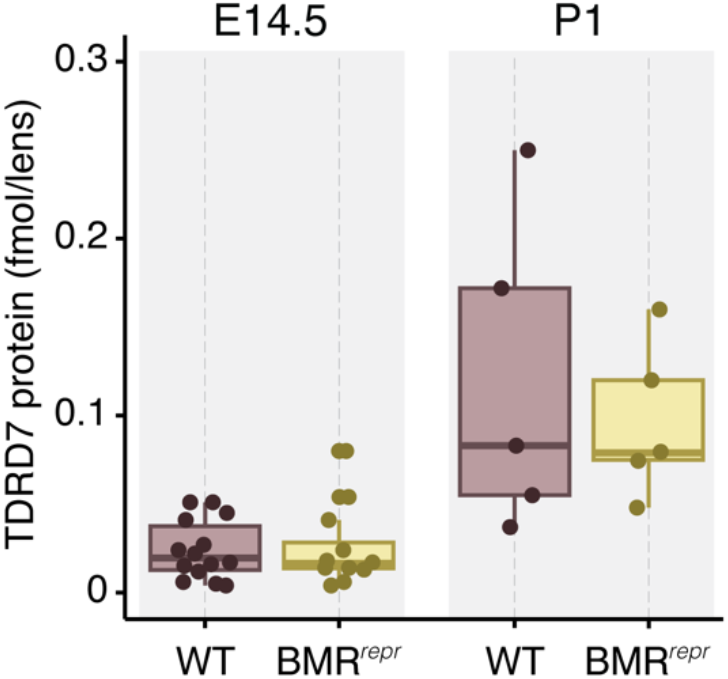
*Tdrd7* expression differences in BMR^*repr*^ mice are buffered at the protein level. Absolute quantification of *Tdrd7* protein in the lenses of E14.5 and P1 wild-type (purple) and BMR^*repr*^ mice (yellow) shows a similar *Tdrd7* protein content.

## DISCUSSION

Comparative genome analysis has revealed many associations between particular traits and genomic changes in particular genes or regulatory elements. Understanding the contribution of mutations in such genes or regulatory elements to the evolution of a trait requires an experimentally-tractable system, such as the mouse, where specific mutations can be introduced and their effects quantified. Gene knockout studies in mice contributed to our understanding of the functional role of many genes naturally lost in a variety of mammalian species (39–46). However, the effect of other, much more frequent genomic changes, such as amino-acid changing mutations in protein-coding sequences and sequence changes in regulatory elements has rarely been explored.

Here, we experimentally investigated the effects of prominent mutations in a lens regulatory element that were observed in the blind mole rat, a burrowing mammal with greatly reduced eyes. To this end, we used CRISPR-Cas9 to precisely replace the mouse sequence of this regulatory element by the BMR sequence. Consistent with our *in vitro* results showing that the BMR sequence exhibits a partial loss of repressor activity in lens cells, we found that BMR^*repr*^ mice exhibit an up-regulation of the *Tdrd7* gene in developing lens. Strikingly, replacing the sequence of this single regulatory element is sufficient to cause a more than two-fold increase in expression, which reveals that this repressor element has a major effect on *Tdrd7* expression. This result contrasts previous findings that perturbations of individual regulatory elements generally have no measurable or only a small effect on gene expression (47–53). Previously-observed robustness at the transcriptional level can be attributed to functional redundancy, whereby multiple regulatory elements control the expression of a gene and a substantial change in gene expression can only be achieved by perturbations of multiple elements (50). Importantly, while expression robustness of other genes may be achieved at the transcriptional level, our observations that the large increase in *Tdrd7* mRNA does not lead to a corresponding increase in protein suggests that robustness of TDRD7 protein expression is instead achieved at the posttranscriptional level. Buffering mechanisms at the protein level are quite frequent, especially for proteins that are part of protein complexes (54). The mechanisms that confer posttranscriptional buffering of increased *Tdrd7* mRNA levels remain to be investigated.

The absence of increased TDRD7 protein levels in the lens likely explains why BMR^*repr*^ mice do not exhibit an obvious lens phenotype. Nevertheless, it is possible that these animals exhibit more subtle phenotypes for which it is hard to detect significant differences with a limited number of individuals. For example, we observed that BMR^*repr*^ mice exhibit slightly more cataract aggregates in the lens; however, many more individuals would be needed to assess whether this subtle difference is consistently observed and statistically significant. In addition, stronger phenotypes may not express under the constant conditions of the artificial laboratory environment. In contrast, in the wild, where environmental fluctuations occur, pathogens exist, and food and mates are limited, both subtle as well as condition-dependent phenotypes would be subjected to natural selection over longer periods of time.

While perturbations or deletions of single regulatory elements can result in drastic (55–57) or subtle phenotypic changes (47,49,52), often they do not result in detectable phenotypic changes (48,50,51,58). Even in the absence of detectable phenotypes when perturbing a single regulatory element, mutations in several regulatory elements can collectively account for phenotypic differences. For example, most mutations associated with complex traits and diseases are located in non-coding, presumably regulatory, genomic regions (59,60). Individually, most of these mutations have a small effect, but collectively they can explain a large portion of the heritability of complex traits (61–64). Similarly, our study showed that perturbing a single lens repressor element in a model organism, where the entire genetic machinery required for functional eyes is intact, did not result in a detectable organismal phenotype. However, we have previously shown that numerous components of the genetic “eye-making” machinery are highly diverged in the BMR and other subterranean mammals, as these species lost several eye-related genes and exhibit divergence in hundreds of eye regulatory elements (21–23). This suggests that eye degeneration in subterranean mammals most likely has a polygenic basis and that perturbations of several regulatory elements are necessary to obtain a detectable phenotype. Ongoing advances in genome engineering (65) will make it easier to test this in future by introducing additional regulatory mutations in the BMR^*repr*^ mouse line. Such studies will likely expand our understanding of regulatory robustness and could ultimately enable predictions of which and how many perturbations are required to cause a particular phenotypic difference.

## METHODS

### Computational identification of *Tdrd7* regulatory locus

Two CNEs upstream of *Tdrd7* were identified in two previous genome-wide screens for CNEs that are diverged in subterranean mammals (21,22). In the first screen, we generated a multiple whole-genome alignment with mouse (mm10 assembly) as the reference and 24 other mammals, used PhastCons (66) to obtain conserved regions, and extracted CNEs by excluding protein-coding regions. This resulted in 351,279 CNEs. For each CNE, we then used the Forward Genomics branch method (13) to assess whether the CNE sequence is preferentially diverged in subterranean mammals (21). In the second screen, we considered the same set of CNEs, but determined whether a CNE exhibits preferential divergence of eye transcription factor binding sites using REforge (22). Applying GERP++ (26), another method to determine evolutionary constraint, to the same multiple genome alignment shows that both PhastCons CNE are part of a larger GERP++ CNE. To determine whether this CNE overlaps previously-published regulatory data, we mapped ChIP-seq, DNaseI-seq, and ATAC-seq data (21,27–32) derived from developing and adult mouse eye tissues to the mouse mm10 assembly using the bedtools suite (67). We inferred TF binding sites using MAST (68) with a previously obtained set of 28 eye TFs (22).

### *In vitro* test for regulatory activity with a dual luciferase reporter assay

We synthesized and cloned a 410 bp mouse sequence that spans both diverged CNEs (mm10 assembly; chr4:45967574-45967983) and the orthologous 208 bp blind mole rat sequence (nanGal1 assembly; KL203377:255659-255866) into a luciferase pGL4.23[luc2/minP] vector (Promega, US). We cloned the insert in the same orientation with respect to the minP promoter and luciferase gene as the CNE and the *Tdrd7* promoter are oriented in the genome. We used the renilla pGL4.73 [hRluc/SV40] (Promega, US) as the control plasmid.

We obtained an aliquot of a primary culture of 21EM15 mouse lens cells for this experiment (kind gift from Dr. Salil Lachke). These cells were cultured in DMEM (Thermo Fisher Scientific, US) supplemented with 10% fetal bovine serum (Merck, Germany), at 37°C, 5% CO2 and 100% humidity. For detachment, cells were washed with PBS and treated with 0.05% Trypsin/EDTA (Thermo Fisher Scientific, US) for 3 minutes at 37°C. After incubation, fresh medium was added, spun down at 140 g for 5 minutes and re-suspended in fresh complete media. 3000 cells/well were seeded into the 384-well plate (Corning, US). The enhancer constructs were transfected 24 h after seeding using FuGENE6 (Promega, US) as the transfection reagent, following manufacturer’s instructions. In brief, FuGENE6 was mixed in OptiMEM (Thermo Fisher Scientific, US) and incubated for 5 min at room temperature. The plasmids with firefly and renilla luciferases were added to the mix in a ratio of 100:1 and, after an incubation period of 20 min at room temperature, the transfection complex was added to the cells in a 4:1 FuGENE/DNA ratio. The luminescence read-out was obtained 48 h after transfection, using the Dual-Glo Luciferase Assay System Kit (Promega, US) as substrate, according to manufacturer’s instructions, and an Envision 2104 Multilabel reader (Perkin Elmer, US) with an ultra-sensitive luminescence 384-well aperture.

We repeated the assay twice (two different plates) with a total of eight technical replicates per plate for each of the mouse and blind mole rat tested sequences. The ratio of firefly and renilla luciferases was normalized by the mean of the replicates of the empty control vector of the respective plate. Significance was assessed using a two-sided Wilcoxon rank sum test.

### Mouse transgenics

#### Ethics statement

Work with mouse was performed in accordance with the German animal welfare legislation and in strict pathogen-free conditions in the animal facility of the Max Planck Institute of Molecular Cell Biology and Genetics, Dresden, Germany. Protocols were approved by the Institutional Animal Welfare Officer (Tierschutzbeauftragter), and necessary licenses were obtained from the regional Ethical Commission for Animal Experimentation (Landesdirektion Sachsen, Dresden, Germany).

##### 1. CRISPR guide design

To measure the molecular and morphological effects of divergence in the *Tdrd7* regulatory locus, we used the CRISPR-Cas9 system to create a transgenic mouse line in which we replaced the above-mentioned 410 bp mouse sequence by the orthologous 208 bp blind mole rat sequence. We designed four guide RNAs to target the *Tdrd7* locus for the CRISPR transgenesis using the Geneious software (8.1.6; Biomatters), and created a repair construct comprising 500 bp homology arms flanking the 208 bp blind mole rat sequence. Guide sequences (PAM sequence underlined) and repair construct sequence (blind mole rat sequence underlined) are the following: gRNA1: 5’GTTCATGGATGGGGGTAGC<u>AGG</u> 3’; gRNA2: 5’ TGATTAATCATTAATGACA<u>TGG</u> 3’; gRNA3: 5’TAATCCCCTGGAGGGTGATC<u>AGG</u> 3’; gRNA4: 5’GATCACCCTCCAGGGGATT<u>AGGG</u> 3’; repairsequence 5’-CAGCAACTATTACAGCCTAATTTTTGTACCAGTAGGTTTATCCATGATTTCAGTGCCTTGTTTGAAGCACCAGTTAGCTAATCTCATGCCTTGCTAAATAGCTGGGCCTCTTTTCTCAGCTTCCTGATAGTGCTTAGTAGACACCATCTGTATTGCAAGTCCTGCCTAAGCTGTATGAATGATGCACAATTTGAAGCAGGAAAGTGAAGATGGTCTCTTAGACAGCAGGTTCTACCCTCTAGGTTCTCTACGCAGAGTGTTCAAACAGTGGTTCTCAACTTATGGGTCTCGACCCCTTTGTCGGGTAAACAATCCTTTTCACAGGAGTCAGATATCCTGCATAACAGGTATTTACATTATGATTAATAGTAGTAGCAAAATCACAGTTATGAAGTAGCAAGAAAAAAATAATTTTATGTCTGGGGGGGTCACTACAACATGAGGAACTGTGTTAAAGCGGTTAAGCTTTACGAAGGTTGAGAGACACTGTGTTAAAGCTT<u>TCATTATCTCCTCTTTCCCATTGTTGGTGGCTGTGTGGCCTCTAAAGCTTGGGTGATCAGTTCCTGCTGGGGATTAGGGCCTTTCAGCTGATTCACCTTAAAGGCCTTTGTGGGCAGCTTAGGGGTTTTGCTGTATCAGCACGTTAACGATTAATTGAAAACAGCTCTGTCCAGAGATCCCTACTTTTCCTGACTTTGTATAAGCAAC</u>CAGTGCCTGAGAAGCTGGGTTCCACCTGGATGCAAAAACAGCTCAGCACAATGGCTGAGTGGCAGGCGGAATGCTGCAAAAGCCACTTGCAGCTTAGCAACTAGGAGAATGGGGACTCAGTTAAG CAGTACACAGGGGGGATGGAAAGCTTCGCCCTCCTCGCACTGCTATTCTGTGAGAATCCTCAGAGGAACAAAGCACCATCTTGAAACTCAATTAAGAGAGCAATAGGTTTGGGGGAAGGGCTGTGAGGGAGGTTTCCAGAGGAGTACCACCCATGACACTGAATAATGTCACAAGCCAGTCCCTGAAGTTCACTGACGGGATGGAAAGTGACAGAAATCCGAGTCCTGTGGATGTGTTTGGGGTTTTCTTTTTGGAGACAATATAAAGAAGCGACAGATGTTCTGGATAGAAGGTGAGATAGCAGCGCTTGAAGTGTCTCCAAATGCCACTTGAGATGTGGAAGGGGGGATTTGCAACCCTTCCCAGTATTTGTG

##### 2. Creation of transgenic embryos

We induced superovulation in female C57Bl6/JCrl donor mice following standard hormonal treatment with PMSG (pregnant mare’s serum gonadotrophin) and HCG (human chorionic gonadotrophin) and performed pronuclear injection of fertilized mouse oocytes following the procedure described in (69). Superovulating females were mated with C57Bl6/JCrl male mice 46 hours between the PMSG and HCG injections at the midpoint of the dark period (12 hours/12 hours, 6am to 6 pm light cycle). After positive plug detection in the morning, the cumulus complexes were isolated and zygotes removed with a treatment of hyaluronidase (final concentration of 0.1% [801 unit/ml]. Before injecting the CRISPR solution, we incubated the recombinant Cas9 protein (ToolGen, 31.31 pmol final) in protein buffer (20 mM Hepes pH 7.5, 150 mM KCl) with 7.8 pmol of each gRNA, 15.6 pmol of tracrRNA, and 0.16 pmol of the gblock repair construct in a total volume of 15 μl, for 15 min at 37°C, allowing the mix to form a functional complex. We next centrifuged the mix (1 min, 13,000 g) through a Durapore PVDF 0.22-μm filter (Merck Millipore, US). We injected the CRISPR-Cas9 solution into the male pronucleus of fertilized zygotes with a motor-driven manipulator-based microinjection stage. Recipient Crl:CD1(ICR) female mice were mated with sterile (vasectomized) Crl:CD1(ICR) males. Around two hours after injections of CRISPR-Cas9 mix into fertilized zygotes, the surviving embryos were transferred into the pseudopregnant recipient female mice (around 20 embryos per recipient) following the procedure described previously (70).

##### 3. Genotyping

We extracted genomic DNA from mouse tails using the QuickExtract DNA extraction kit (Epicentre) following manufacturer’s instructions, and performed PCR using the Phusion Flash High-Fidelity PCR master mix (Thermo Fisher Scientific, US). Two primer pairs were designed to bind to the 5’ and 3’ flanks of the mouse sequence outside of the homology arms and to the blind mole rat specific sequence. PCR products were Sanger-sequenced to confirm the precise replacement of the regulatory element. Primer sequences are the following: 5’ flanking_forward (mouse): 5’TCAGTCTGTGAGCCGTTGAC 3’; 5’ flanking_reverse (BMR^*repr*^ mouse): 5’TAACGTGCTGATACAGCAAAACC 3’; 3’ flanking_forward (BMR^*repr*^ mouse): 5’GCTTAGGGGTTTTGCTGTATCA 3’; 3’ flanking_reverse (mouse): 5’ACACAACCCAAGATCCACACG 3’.

### Quantification of *Tdrd7* mRNA levels by RT-qPCR

We quantified *Tdrd7* mRNA expression by RT-qPCR, with primers to amplify 97 bp of the mRNA spanning exons 12 and 13. We used the ribosomal *Rpl13a* gene as control. Primer sequences are the following: Tdrd7-fw: 5’TGGCAATTCGACATCCGTGA 3’; Tdrd7-rev: 5’ GCACTTTATAGCCTGAGGGGG 3’; Rpl13a-fw: 5’ CCCACAAGACCAAGAGAGGC 3’; Rpl13a-rev:5’ CACCATCCGCTTTTTCTTGTCA 3’. We dissected lenses and testes/ovaries of E14.5, P1, and 12 week-old adult female and male wild-type and homozygous BMR^*repr*^ mice and snap-froze collected tissues in liquid nitrogen. Total RNA was extracted with a standard TRIzol RNA extraction and chloroform/isopropanol precipitation and cDNA was prepared with the ProtoScript II First Strand cDNA synthesis kit (New England Biolabs, Frankfurt, Germany). For each sample, we collected at least three biological replicates, and measured each at least 3 times (technical replicates) in the LightCycler96 (Roche, Germany). We detected expression using SYBRgreen marker (Roche, Germany). To obtain the normalized expression values of *Tdrd7* in BMR^*repr*^ samples with respect to the wild-type condition, we followed the procedure described in (71). In short, we computed the ΔCq of each BMR^*repr*^ sample with respect to the average of wild-type samples, corrected those for primer efficiency, and normalized by the control gene *Rpl13a*. We tested the difference in normalized *Tdrd7* expression between wild-type and BMR^*repr*^ mice with a two-sided T-test.

### Absolute quantification of *Tdrd7* protein by MS Western

We opted for an absolute quantification of *Tdrd7* protein in developing lenses using MS-Western (72). *Tdrd7* was quantified by co-digestion of its SDS PAGE-separated band with the band of isotopically labeled chimera composed of concatenated proteotypic peptides.

#### 1. Preparation of labeled *Tdrd7* peptides

We selected quantotypic peptides for *Tdrd7* protein using the collection of mass spectra available in the MaxQB database (Supplementary Table 1), according to (73). Peptide sequences were included into an E. *coli*-optimized synthetic gene construct containing a TwinStrep tag, selected phosB and BSA peptides (72). The construct was cloned into a pET expression vector and transformed into an BL21 (DE3) E. *coli* strain, auxotrophic for arginine and lysine. Cells were grown at 37°C in MDAG-135 media (74), supplemented with 16 unlabeled amino acids and isotopically labeled ^13^C_6_^15^N_4_-l-arginine and ^13^C_6_-l-lysine (Silantes, Munich, Germany). Protein expression was induced by 1 mM isopropyl β-d-1-thiogalactopyranoside (IPTG) for 3 hours. After induction, cells were pelleted, re-suspended in 2x phosphate-buffered saline (PBS), snap-frozen in liquid nitrogen, and stored at −80°C. Prior to analyses, cells were thawed and lysed in an equal volume of 2x Laemmli buffer by incubating at 95°C for 10 minutes. The sample was clarified by centrifugation and the supernatant subjected to 1D SDS PAGE on a 4-20% pre-cast gradient gel. Proteins were visualized by Coomassie staining.

#### 2. Sample preparation

We carefully dissected the lenses of wild-type and homozygous BMR^*repr*^ E14.5 embryos and P1 pups (2 biological replicates for E14.5 samples and a single replicate for P1), snap-froze them in liquid nitrogen and stored at −80°C until use. To lyse the tissue, we added 40μl of 1x Laemmli buffer with 0.5μl of protease inhibitors cocktail (Thermo Fischer Scientific, US) and 20-30 0.2mm stainless steel beads (Next Advance, US) per vial, and homogenized it in a TissueLyser (Qiagen, Düsseldorf, Germany) for 10min at 4°C. Homogenized material was centrifuged for 10 minutes and heated at 80°C for 5 minutes. After cooling down to room temperature, cell debris were spun down and the supernatant loaded onto a 1D SDS min-gel (BioRad Laboratories, US). We visualized the gel by Coomassie staining and excised the region corresponding to the molecular weight of *Tdrd7* protein for subsequent MS-Western analysis.

#### 3. Mass spectrometric analysis and absolute quantification using MS-Western

Absolute quantification of *Tdrd7* protein was performed as described in (72). Briefly, an aliquot of 0.5 pmol of BSA standard (Thermo Fischer Scientific, US) was analysed by 1D SDS PAGE, visualised by Coomassie, and the BSA band was excised. The gel slices containing TDRD7, the isotopically labelled chimeric protein standard from a separate gel, and a band of the BSA standard were combined together. Proteins were reduced with 10mM DTT, alkylated by iodiacetamide and in-gel digested overnight with trypsin. The resulting peptide mixture was extracted twice by exchange of 5% formic acid and acetonitrile, extracts pooled together and dried in a vacuum centrifuge. Peptides were re-suspended in 25μl of 5% FA and a 5 μl aliquot was analysed by LC-MS/MS on a nano-UPLC system interfaced to a Q Exactive HF Orbitrap mass spectrometer (both Thermo Fischer Scientific, Bremen, Germany). The nano-UPLC was equipped with an Acclaim PepMap100 C18 75 μm i.d. × 20 mm trap column and 75 μm × 15 cm analytical column (3μm/100A, Thermo Fisher Scientific, Bremen, Germany). Peptides were separated using an 80-minute linear gradient: solvent A was 0.1% aqueous formic acid and solvent B was 0.1% formic acid in neat acetonitrile. Three blank runs were performed after each sample analysis to minimize carryover, the last blank run was recorded and searched against mouse sequences in UniProt database. To set up PRM parameters, the relevant information on target peptides (m/z, charge state, retention time; Supplementary Figure 1) was obtained in preliminary experiments using a tryptic digest of the chimeric standard. In all experiments, FT MS spectrum was acquired within the mass range of m/z 350-1700 at the mass resolution of R_m/z=200_ = 240000 (FWHM); targetAGC of 3×10^6^, 150 ms maximum injection time. It was followed by PRM scans at R_m/z=200_ = 120000 resolution; target AGC of 1×10^5^; 200 ms maximum injection time triggered by a scheduled inclusion list. Lock mass was set to the singly charged ion of dodecamethylcyclohexasiloxane ion ((Si(CH3)2O))6; m/z = 445.1200).

We processed the spectra using the SkyLine software (75). We manually verified the data, refining the peak boundaries where necessarily. MS/MS spectra from PRM experiments were also searched with MASCOT software against a database of mouse protein sequences. Quantification was performed using the sum of extracted peak areas of 2-4 most abundant y-ion fragments whose m/z exceeded the m/z of the corresponding precursor ion and integral value exceeding 1×10^6^. First, the amount of each labelled standard peptide was quantified using the peak areas of peptides of unlabelled BSA standard (note that all peptides in chimera standard protein, including peptides from BSA and TDRD7, were present in equimolar ratios). Then, the molar abundance of the labelled standard peptides was used to quantify peptides from endogenous TDRD7. As data quality control, we required that normalized ratios of fragment ion abundances were the same in labelled (from chimera) and corresponding unlabelled (endogenous) peptides. Finally, the data were normalized to the number of eye lenses pooled in each sample.

### Analyzing lens morphology

#### 1. Quantifying nuclear cataracts

To quantify the number and size of cataract aggregates in the lenses of adult mice, we imaged them with an Olympus SZX16 stereoscope equipped with a q-Imaging camera. We carefully dissected the lenses of wild-type and homozygous BMR^*repr*^ mice with ages between 6 months and 1.5 years, immediately embedded them in Optiprep refractive index matching media (76), and imaged under darkfield illumination (11.5x magnification; 250 ms exposure and 6.5 gain). We used FIJI (77) to process images with a script that includes subtracting background (rolling ball radius = 50 pixels), median filtering (radius = 2 pixels), detecting objects with the Interactive Watershed plugin (SCF; seed dynamics 2000, intensity threshold 4000, peak flooding 50, AllowSplitting = false), and finally obtaining the number and size of elements with Analyze Particles plugin (size = 1000-10000 px^2^).

#### 2. Scanning electron microscopy

Lenses of 2 year old wild-type and homozygous BMR^*repr*^ mice aged were carefully dissected and fixed in a modified Karnovslky’s fixative (2% glutaraldehyde/ 2% formaldehyde in 100 mM phosphate buffer) at 4°C for at least one day to allow the fixative to penetrate through the more external layers of the lens. After this period, the lenses were cut in half through the equatorial plane and fixed again in the same solution for at least another 3 days. Samples were washed with PBS (10 mM Na_2_HPO_4_, 1.8 mM KH_2_PO_4_, 137 mM NaCl, 2.7 mM KCl, pH 7.4) for 5x 5min. After that, the material was postfixed in 1% osmium tetraoxide in PBS, washed several times in PBS and water, and dehydrated in a graded series of ethanol/water mixtures up to pure ethanol (30%, 50%, 70%, 96% and 3 × 100% on molecular sieve, 15 minutes each). Samples were critical point dried using a Leica CPD300 (Leica Microsystems, Wetzlar, Germany) and mounted on 12 mm aluminium stubs using conductive carbon tabs. The lens halves were additionally grounded with conductive liquid silver paint. To increase contrast and conductivity, the samples were sputter coated with gold (BAL-TEC SCD 050 sputter coater, settings: 60 sec, with 60 mA, at 5 cm working distance). Finally, the samples were imaged with a JSM 7500F scanning electron microscope (JEOL, Garching, Germany) running at 5 kV (lower SE-detector, working distance 8 mm).

## Acknowledgment

We thank Maurício de Rocha Martins for helpful discussions and comments on the manuscript, Salil Lachke for providing the lens cell line, Andrea Schuhmann for assistance in MS experiments and analyses, and the following MPI-CBG facilities for their support: Antibody Facility, Biomedical services, Cell Technologies Group, DNA sequencing, Mass Spectrometry, Transgenic Core Facility, TransgeneOmics, and Technology Development Studio. This work was supported by the Max Planck Society.

## Supplementary materials

**Supplementary Table 1.**
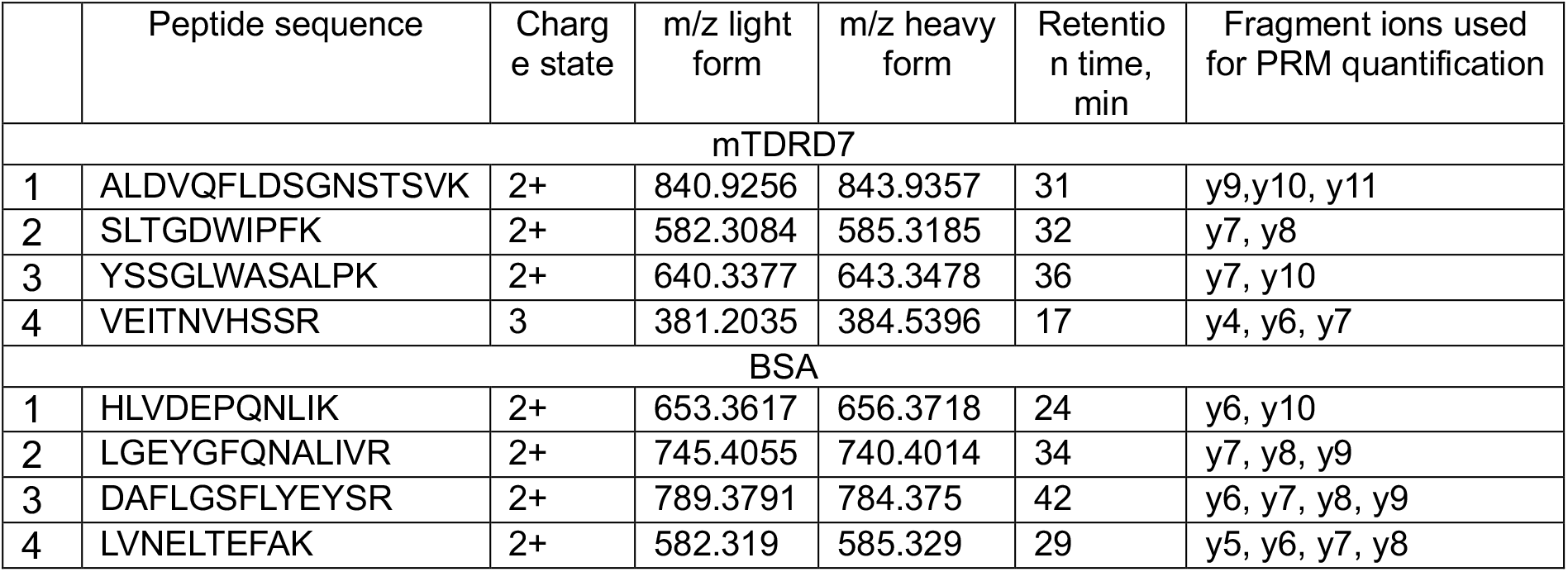
Description of quantotypic peptides selected for the absolute quantification of TDRD7 protein.

|  | Peptide sequence | Charge state | m/z light form | m/z heavy form | Retention time, min | Fragment ions used for PRM quantification |
| --- | --- | --- | --- | --- | --- | --- |
| mTDRD7 |  |  |  |  |  |  |
| 1 | ALDVQFLDSGNSTSVK | 2+ | 840.9256 | 843.9357 | 31 | y9,y10, y11 |
| 2 | SLTGDWIPFK | 2+ | 582.3084 | 585.3185 | 32 | y7, y8 |
| 3 | YSSGLWASALPK | 2+ | 640.3377 | 643.3478 | 36 | y7, y10 |
| 4 | VEITNVHSSR | 3 | 381.2035 | 384.5396 | 17 | y4, y6, y7 |
| BSA |  |  |  |  |  |  |
| 1 | HLVDEPQNLIK | 2+ | 653.3617 | 656.3718 | 24 | y6, y10 |
| 2 | LGEYGFQNALIVR | 2+ | 745.4055 | 740.4014 | 34 | y7, y8, y9 |
| 3 | DAFLGSFLYEYSR | 2+ | 789.3791 | 784.375 | 42 | y6, y7, y8, y9 |
| 4 | LVNELTEFAK | 2+ | 582.319 | 585.329 | 29 | y5, y6, y7, y8 |

**Supplementary Figure 1.**
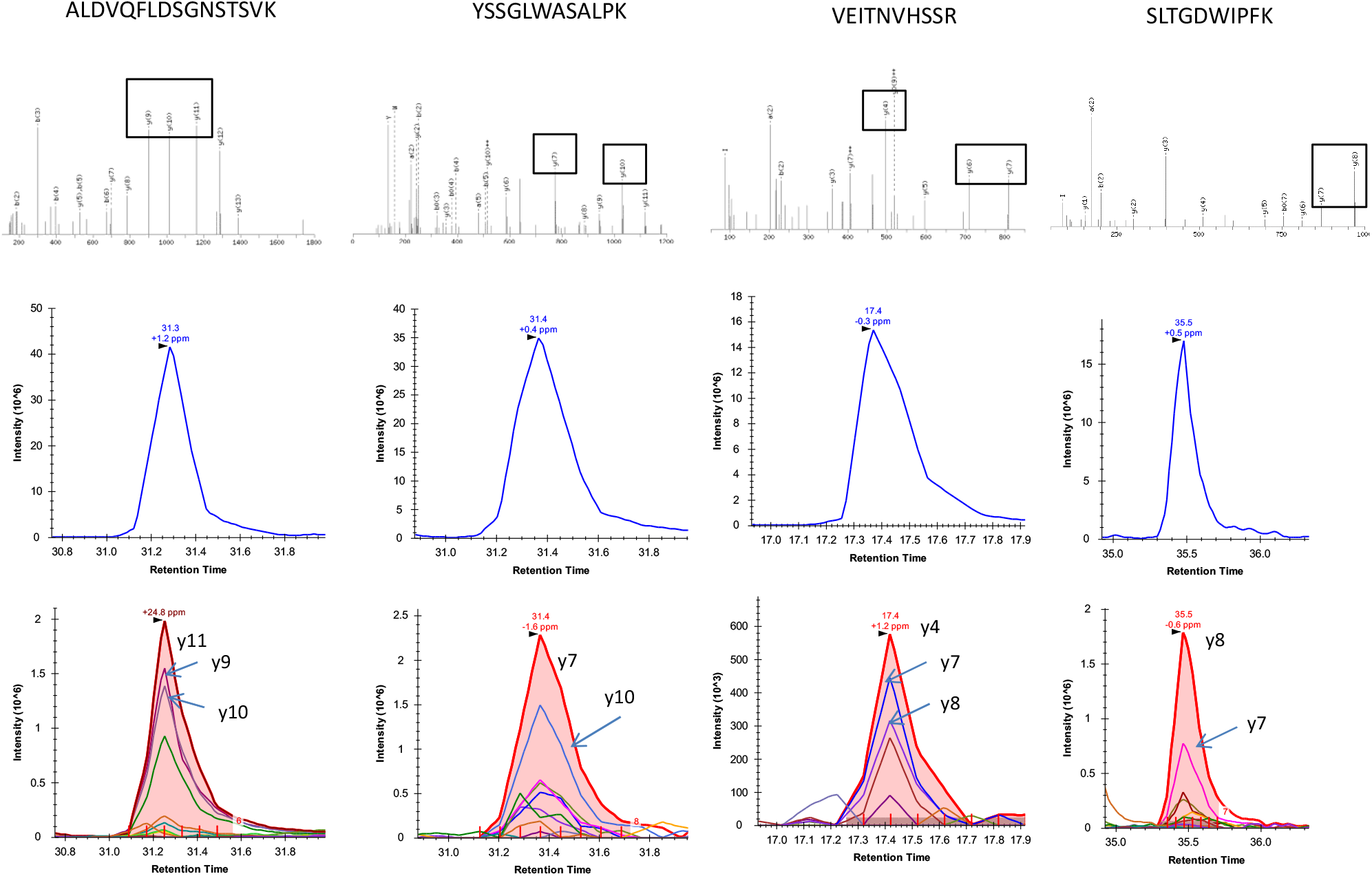
Alignment of XIC profiles of *Tdrd7* quantotypic peptide standards. Four tryptic peptides were selected from the *Tdrd7* sequence for its absolute quantification. Upper row: MS/MS spectra of 15N, 13C-labelled peptide standards; fragments selected for the quantification are boxed. Their XIC are shown below.

